# Impaired Complex I dysregulates neural/glial precursors and corpus callosum development revealing postnatal defects in Leigh Syndrome mice

**DOI:** 10.1101/2025.05.16.654318

**Authors:** Sahitya Ranjan Biswas, Porter L. Tomsick, Colin Kelly, Brooke A. Lester, Julia P. Milner, Sara N. Henry, Yaris Soto, Samantha Brindley, Nicole DeFoor, Paul D. Morton, Alicia M. Pickrell

**Affiliations:** Translational Biology, Medicine, and Health Graduate Program, Virginia Polytechnic Institute and State University, Roanoke, VA 24016 USA; School of Neuroscience, Virginia Polytechnic Institute and State University, Blacksburg, VA 24061 USA; Department of Biomedical Sciences and Pathobiology, Virginia-Maryland College of Veterinary Medicine, Virginia Polytechnic Institute and State University, Blacksburg, VA 24061 USA

**Author notes:** Correspondence should be addressed to: Alicia M. Pickrell, Life Science I Room 217, 970 Washington Street SW, Blacksburg, VA 24061 USA; Paul D. Morton, Life Science I Room 223, 970 Washington Street SW, Blacksburg, VA 24061 USA.

**Keywords:** Leigh syndrome, Complex I, postnatal neurogenesis, neural stem cells, subventricular zone, corpus callosum, mitochondria, oligodendrocytes, neurodevelopment

## Abstract

Leigh syndrome (LS) is a complex, genetic mitochondrial disorder defined by neurodegenerative phenotypes with pediatric manifestation. However, recent clinical studies report behavioral phenotypes in human LS patients that are more reminiscent of neurodevelopmental delays. To determine if disruptions in epochs of rapid brain growth during infancy precede the hallmark brain lesions that arise during childhood, we evaluated neural and glial precursor cellular dynamics in a mouse model of LS. Single cell RNA sequencing along with histological and anatomical assessments were performed in NDUFS4 KO mice and compared with controls to determine the impact of Complex I deficiency on neural stem cells, their neuronal and oligodendroglial progeny, lineage progression, and overt differences in specific brain regions. Our findings show disruptions in all categories, specifically within the subventricular zone and corpus callosum. Given that LS is purely considered a neurodegenerative disease, we propose that mitochondrial dysfunction is a neurodevelopmental signature predating classic diagnosis in LS.

## Introduction

Leigh syndrome (LS) is a severe and prevalent inherited primary mitochondrial disorder affecting children, characterized by progressive neurodegeneration and early mortality^1,2^. To date, mutations in more than 110 nuclear or mitochondrial genes have been found associated with the disease, most of which affect assembly subunits or core components of the five complexes responsible for oxidative phosphorylation (OXPHOS)^2,3^. Consequently, the prevailing hypothesis regarding disease pathogenesis is that dysfunctional electron transport chains place increased metabolic stress on high energy-demanding neurons, ultimately leading to their degeneration^4,5^. This neuronal loss is reflected in the hallmark lesions observed in different parts of the brain, including the olfactory bulb, brain stem, cerebellum, basal ganglia, midbrain, pons and thalamus, which also account for the phenotypes of the disease and respiratory distress leading to mortality^6-8^.

LS symptomology has been associated with neurodegeneration, where the loss of previously acquired skills in these pediatric patients has long been considered the most prominent feature of the disease^2^. However, recent follow-up studies in large collective cohorts of LS patients reported that delays in achieving early developmental milestones - such as head movement, rolling, independent ambulation, sitting without support - are often the most common feature, emerging prior to the onset of regressive phenotypes^9,10^. Most research focuses on understanding the neurodegenerative and neuroinflammatory aspects of the disease. But considering that these genetic mutations are inherited, and recent studies indicate that neurodevelopmental defects may occur prior to the onset of neurodegenerative lesions, a more thorough evaluation at the cellular level of the brain, prior to neuron loss, is warranted.

To understand whether key developmental programs underlying early life neurological milestones are affected in LS, we utilized the NDUFS4 knockout (KO) mouse model, a well-established preclinical model of LS characterized by impaired mitochondrial complex I activity^11,12^. This model closely replicates the neuropathological and motor symptoms observed in human patients, including characteristic brain lesions, ataxia, hypotonia, and respiratory failure^13^. Moreover, complex I deficiency remains the most frequent cause of LS, accounting for about one-third of all cases^14^.

Using this model, we identified defects in neural stem and progenitor cell proliferation, a preference for quiescence within the stem cell pool, and a decline in neural progenitors undergoing lineage progression into immature neurons during early postnatal development in the subventricular zone (SVZ). In addition, we found evidence of significant declines in gliogenesis, particularly within the oligodendrocyte lineage, paired with a reduced capacity for myelination of the corpus callosum. Together, our findings indicate a selective vulnerability of neural and oligodendrocyte precursors, resulting in delays in maturation and an underdeveloped corpus callosum prior to the neurodegenerative onset in a LS mouse model. In addition, our transcriptomic data indicate diverse changes amongst many cell populations comprising the neural and glial lineages at early stages of development when there is overlap in neurogenesis and gliogenesis, offering aid in uncoupling the two along with insights into neonatal contributions to the array of developmental delays exhibited in LS patients.

## Results

### Gross neuroanatomical deficits are notable prior to disease onset in NDUFS4 KO mice

Patients with Leigh syndrome are typically diagnosed following the onset of symptoms related to psychomotor decline, which coincide with neurodegenerative lesions found on MRI^15^. Previous evaluation of NDUFS4 KO mice characterized signs of neurodegeneration appearing at postnatal day 37 (P37)^11-13^. To investigate potential early indicators of impaired brain development, we assessed gross brain anatomy that preceded this time point on postnatal days 14, 24, and 30. At these time points, both WT and NDUFS4 KO mice were able to maintain body weight/growth but gains in brain weight lagged significantly **(Figure 1A-B)**. Congruent with differences in gross brain weight, we observed a notable reduction in hemispheric width in NDUFS4 KO mice, particularly at P14 **(Figure 1C-D)**. These data warrant further investigation as they indicate that neurodevelopmental delays precede the documented neurodegeneration in this NDUFS4 KO model of LS.

**Figure 1.**
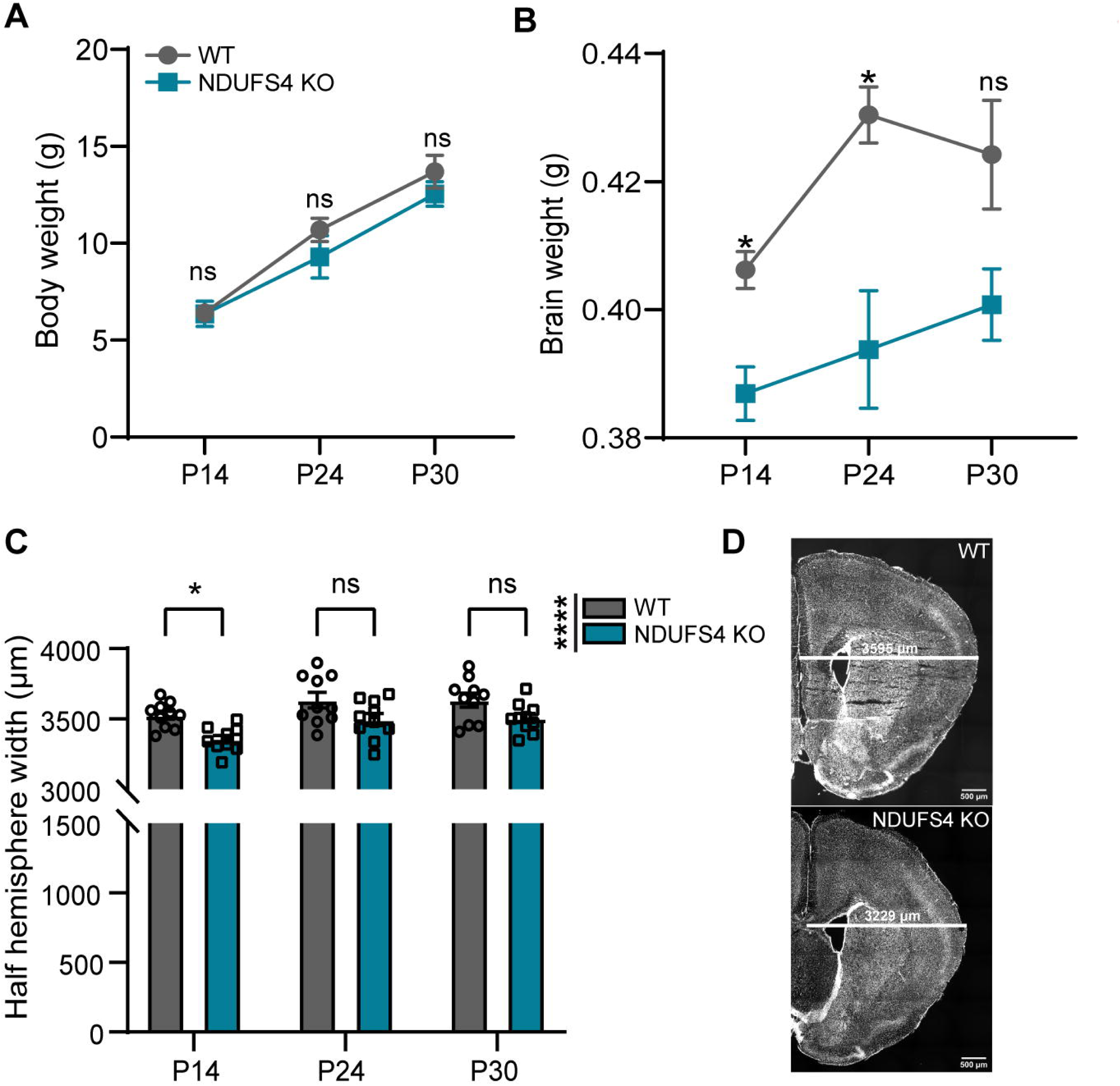
NDUFS4 KO mice exhibit early postnatal alterations specific to brain development. (**A-B).** Overall body weight (A) and brain weight (B) comparisons at P14, 24, and 30 between WT and NDUFS4 KO mice. **C.** Half hemisphere width mean measured between Bregma +1.53- and +1.23-mm. **D.** Representative images from Bregma +1.23 mm of P14 mice. Scale bar= 500 µm. White line denotes where measurements across hemispheres were taken. Two-way ANOVA was performed; *p<0.05, **p<0.01, ****p<0.0001, ns = not significant. Bar plot represents mean and standard error of mean. Each dot represents one animal. n=5/group.

### Reduced neural stem progenitor populations and impaired lineage progression in the SVZ of NDUFS4 KO mice

We next decided to investigate the cell populations that reside in the SVZ, as it remains postnatally active in rodents and higher order species^16-19^. To examine cellular changes in the neurogenic pool of the SVZ, we performed immunohistochemistry (IHC) using SOX2 and doublecortin (DCX) to delineate neural stem/progenitor cells (NSPCs) and neuroblasts, respectively, at 3 postnatal time points preceding neurodegenerative onset. To assess regional differences within the SVZ, we sampled 5 regions on the coronal plane spanning the frontal lobe corresponding with Bregma areas +1.53 to +0.33 mm **(Supplementary Fig. 1A)**. At P14, we observed a significant reduction in the density of SOX2^+^ cells in the NDUFS4 KO indicating a decrease in NSPC numbers; however, these differences were absent at later time points **(Figure 2A-B)**. At this age, we found the more caudal regions of the SVZ accounted for a significant majority of the reduction in NSPC numbers **(Supplementary Fig. 1B)**. We next evaluated lineage progression as SOX2^+^ NSPCs upregulate DCX in transition to neuroblasts. At P14, we found no differences in the number of SOX2^+^ DCX^+^ cells nor in the percentage (SOX2^+^ DCX^+^/SOX2^+^) of NSPCs undergoing transition to the neuronal lineage; however, there was a significant reduction as early as P24 which persisted into P30 in the SVZ of NDUFS4 KOs **(Figure 2C-E)**. This significant reduction was accounted for in the second most rostral area (Bregma +1.23) assessed **(Supplementary Fig. 1C-D)**. We next evaluated neuroblasts and found a significant reduction in the number of DCX^+^ cells at P14 which was not sustained at the later ages assessed **(Figure 2F)**. When evaluating the rostrocaudal distribution of the SVZ, we found that most of this difference was within the more rostral regions assessed **(Supplementary Fig. 1E)**; whereas disruption in lineage progression is evident at later ages, which may indicate NSPCs are shifting toward a less active phenotype or diverging from the neuronal lineage.

**Figure 2.**
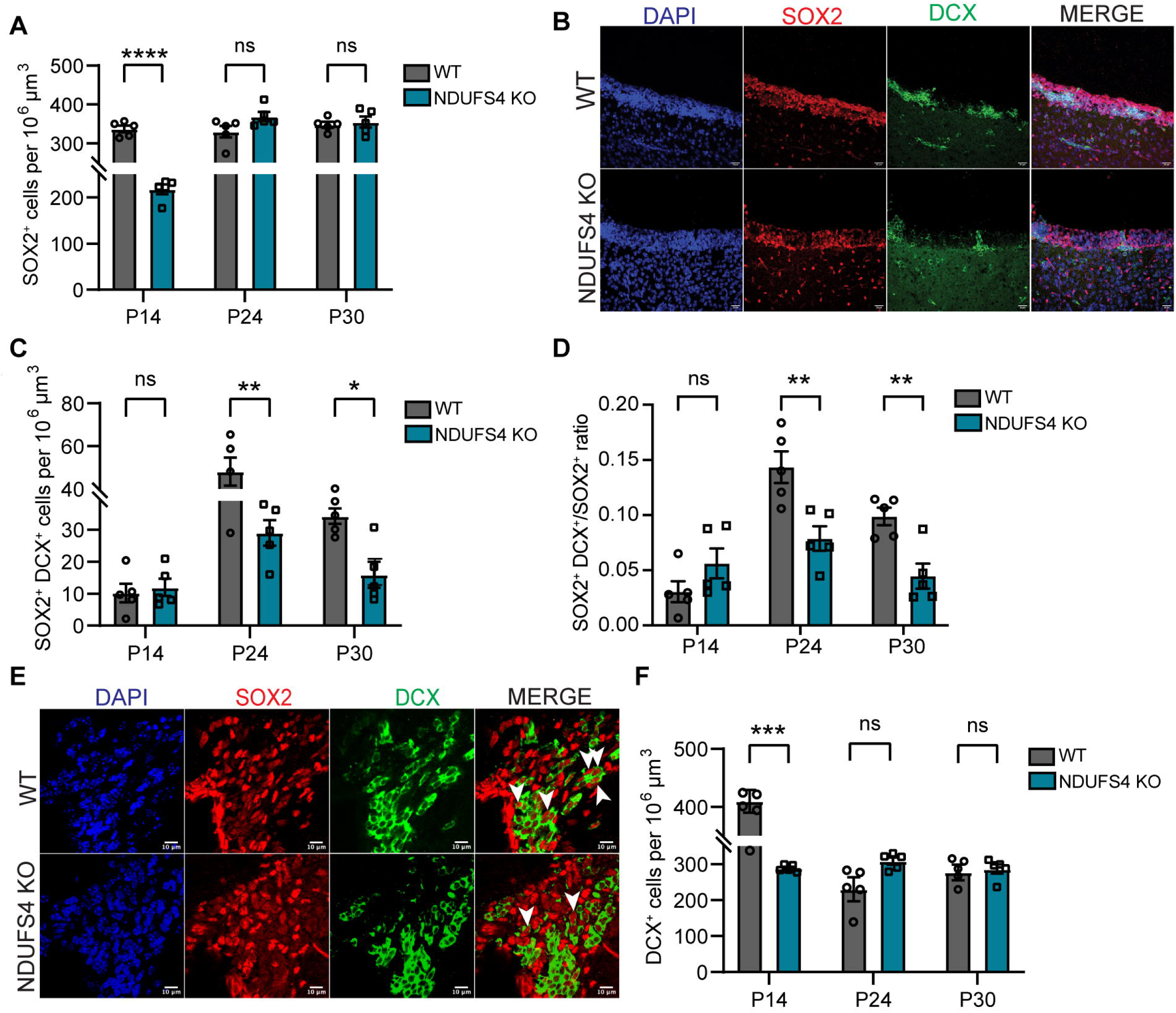
The SVZ displays reduced number of neural progenitors, neuroblast density, and neural stem cell commitment in NDUFS4 KO mice. **A.** Density of SOX2^+^ in the SVZ at P14, P24, and P30. **B.** Representative confocal images of immunocytochemical detection of SOX2 (red), DCX (green), and DAPI (blue) of lateral SVZ wall at P14 in WT and NDUFS4 KO mice. Scale bar= 20 µm. **C-D.** Density of (C) double positive cells (SOX2^+^/DCX^+^) and (D) double positive (SOX2^+^/DCX^+^) /SOX2 ratio at P14, P24, and P30 between groups. **E.** Representative confocal images of immunocytochemical detection of SOX2 (red), DCX (green), and DAPI (blue) of lateral SVZ wall at P24 between WT and NDUFS4 KO mice. White arrows indicate double positive cells. Scale bar= 10 µm. **F.** DCX^+^ cells in SVZ at P14, P24, and P30. Two-way ANOVA was performed; *p<0.05, **p<0.01, ***p<0.001, and ****p<.0001. Bar plot represents mean and standard error of mean. Each dot represents one animal. n=5/group.

Together, these results suggest a biphasic disruption in neurogenesis in complex I deficient mice. At P14, there were fewer neural stem/progenitor cells **(Figure 2)** leading to a reduced generation of neuroblasts, which may in part account for the maturational delays in brain growth exhibited **(Figure 1)**. The reciprocal relationship between reduced NSPCs in the more caudal domains and reduced numbers of neuroblasts in the rostral domains of the SVZ at the earliest age assessed (P14) may also be indicative of reduced tangential migration, owing to fewer numbers of NSPCs generating neuroblasts (**Supplementary Fig. 1**). Considering that mitochondrial disease has genetic origins, we expect that these findings are mainly governed by intrinsic defects in the NSPC pool rather than extrinsic factors such as microenvironmental cues. However, it is possible that other cells in the brain that lack NDUFS4 contribute indirectly.

### Decreased neurogenesis in NDUFS4 KO

While a reduction in NSPC numbers within the early postnatal SVZ may be indicative of cell death, our longitudinal findings demonstrate normal numbers of NSPCs at the expense of a decline in lineage progression **(Supplementary Fig. 1**, **Figure 2);** therefore, we reasoned that these early losses may be due to disruptions in cellular genesis. We next assessed NSPC proliferation within the SVZ of WT and NDUFS KO animals following immunohistochemistry at P14. We found a significant reduction in general cell proliferation within the SVZ following quantification of Ki67^+^ cells **(Figure 3A)**. Regardless of genotype, the average number of proliferating NSPCs (SOX2^+^Ki67^+^) represented 55-57% of cells undergoing mitosis (Ki67^+^) within the SVZ **(Figure 3B)**. However, we found a significant reduction in the proportion of NSPCs undergoing cell proliferation in NDUFS4 KO **(Figure 3C-D)**. Together these findings suggest that complex I deficiency impairs NSPC production which, paired with our earlier findings, contributes to a delay/loss in newborn neurons during a critical postnatal period of brain development.

**Figure 3.**
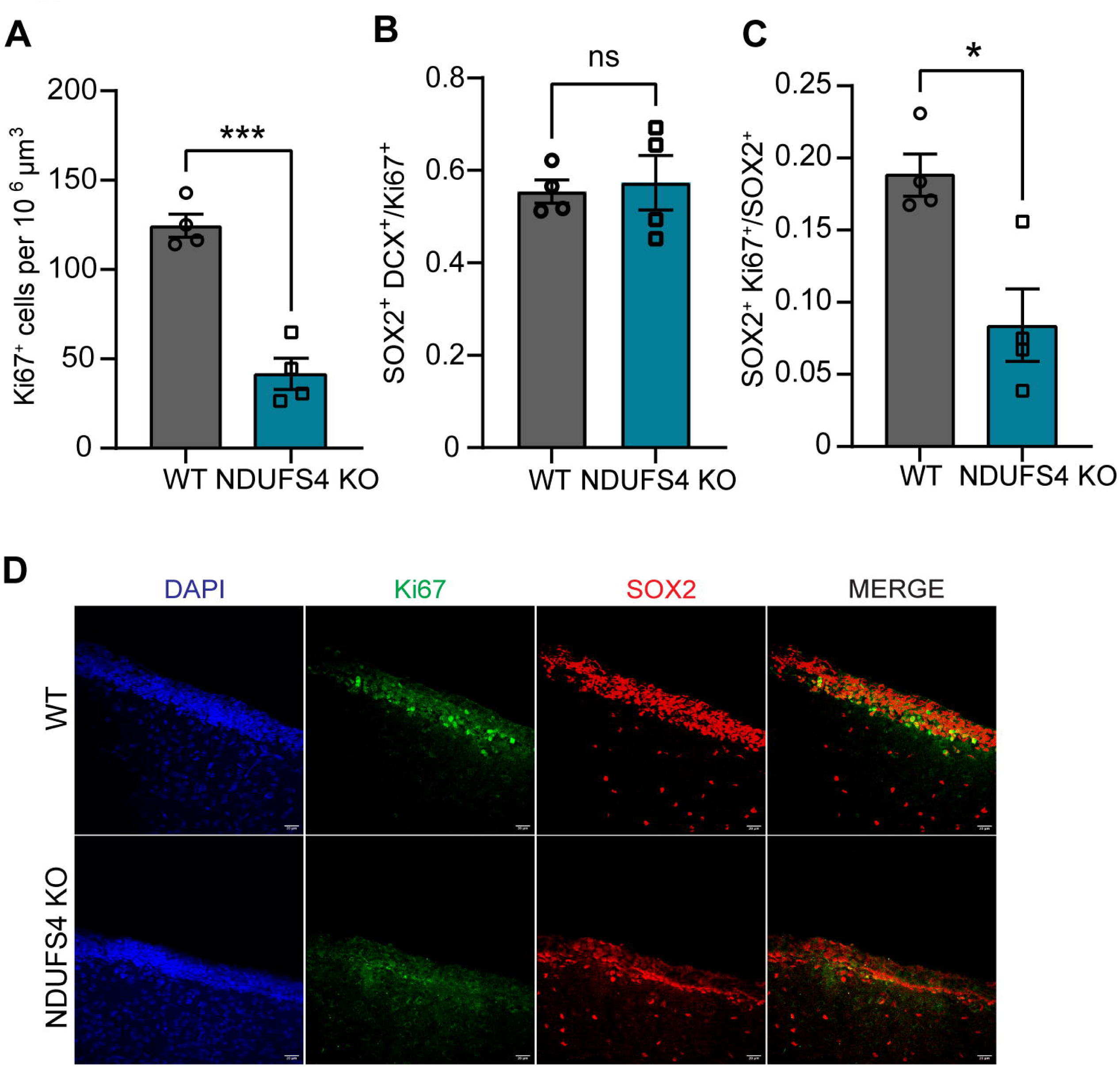
Reduced Proliferation of Neural Progenitors in the SVZ of NDUFS4 KO mice. **A.** Density of Ki67^+^ cells in the P14 SVZ. **B.** Percentage of SOX2^+^ Ki67^+^/Ki67^+^ cells in P14 SVZ **C**. SOX2^+^ Ki67^+^/ SOX2^+^ ratio in the P14 SVZ. **D.** Representative images from lateral ventricles of P14 mice stained for SOX2 (red), Ki67(green), and DAPI (blue). Scale bar = 20 µm. Unpaired t-test was performed; *p<0.05, ***p<0.001. Each dot represents one animal. n=4/group

To corroborate our *in vivo* findings, we performed neurosphere assays to eliminate the influence of systemic and multicellular alterations on NSPC dynamics. Since our most striking findings were at P14, we focused all subsequent analyses on this age. NSCs were isolated from the SVZ of WT and NDUFS4 KO animals at 14 days of age and cultured in the same conditions to assess neurogenesis. Considering that NDUFS4 KO had fewer NSCs to begin with at this age, we plated and quantified the properties of secondary spheres after one passage to ensure we obtained the same starting number of cells prior to analysis. However, neurospheres derived from KO mice were still noticeably smaller and in lower numbers; quantification of sphere diameter from the spheres that were present conferred a significant decrease in size, illustrating a reduction in the capacity to self-renew **(Supplementary Fig. 2)**. When evaluating the distribution of sphere size, there was a significant shift towards smaller spheres, further supporting the notion that mitochondrial function is essential for postnatal production of NSPCs **(Supplementary Fig. 2A)**.Taken together, the recapitulation of our *in vivo* proliferation findings **(Figure 3)** with the *in vitro* neurosphere assay suggests that disruption of NSPC production and lineage progression is governed in part by cell intrinsic mechanisms.

### Single cell transcriptomic profiling of postnatal SVZ identifies distinct classes and subclasses along the neuronal lineage affected in NDUFS4 KO mice

To understand how NDUFS4 impacts neurogenesis within the SVZ, which is occupied by numerous cell types with high transcriptional diversity in the postnatal brain, we performed single cell transcriptomic profiling to better elucidate which biological processes were most affected in its absence. To best align nomenclature, we use the terms neural progenitor cells (NPCs), transit amplifying cells (TAPs), and Type C cells interchangeably. We collected 40,324 cells with a median of 1,223 genes and 1,842 unique molecular identifiers (UMIs) quantified per cell from WT and NDUFS4 KO from microdissected SVZ samples **(Figure 4A).** For higher data resolution, we analyzed cells from all replicates, accounting for batch effects using Harmony. Initial analysis identified 19 distinct cell clusters **(Figure 4A)**, which were assigned to 14 known cell types based on the detection of known marker genes^20^ **(Figure 4A-B)**.

**Figure 4.**
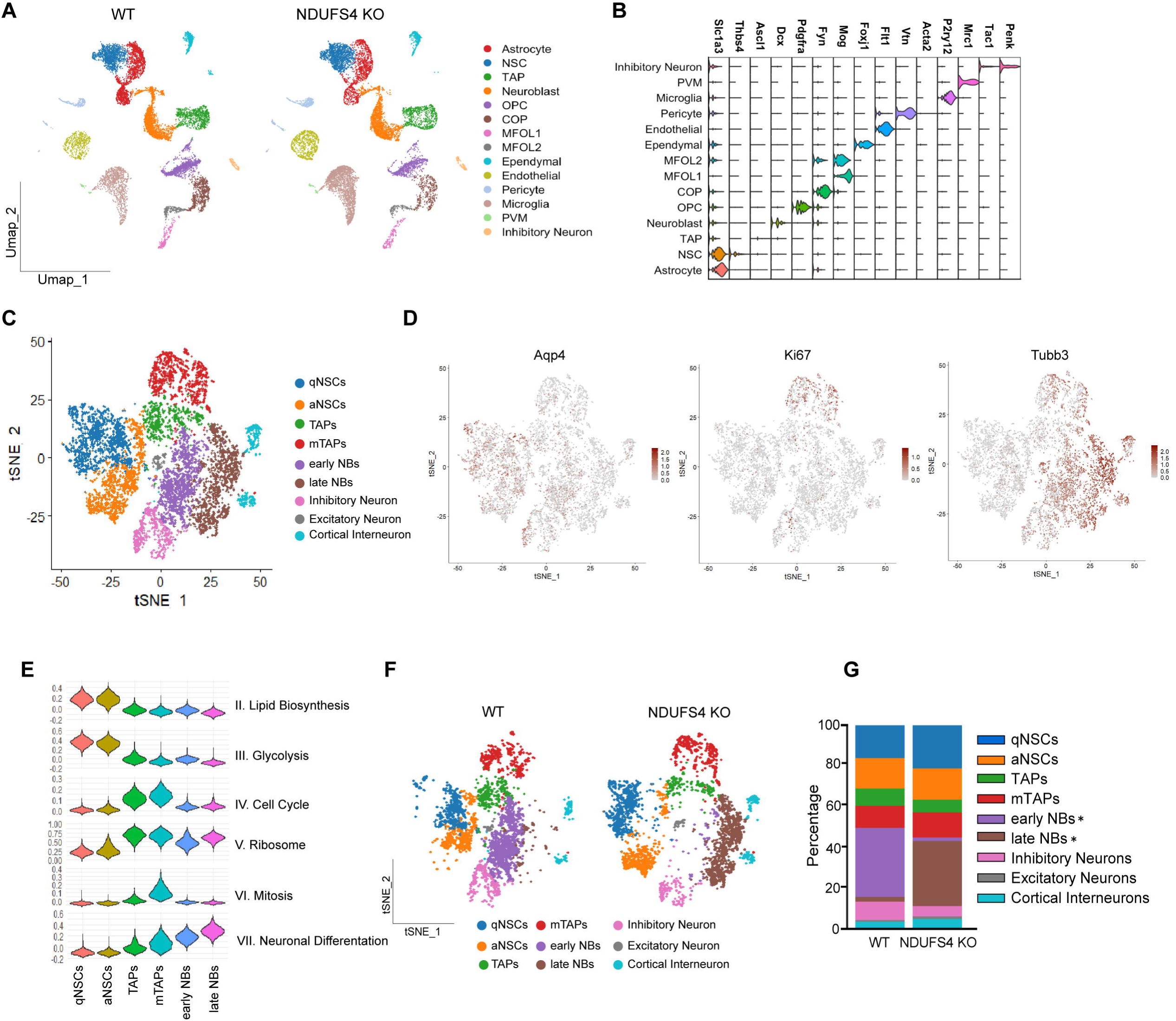
Characterization of cell types and subtypes residing in the P14 SVZ following single-cell sequencing. **A.** UMAP plots of WT and NDUFS4 KO cells colored by cluster annotation. **B.** Expression of known marker genes for each cluster. **C.** tSNE plot for subclusters deriving from NSCs, TAPs, and NBs. **D.** Expression of known markers of different subclusters. **E.** Subcluster characterization based on published gene sets in ^22^ for relevant biological processes. **F.** tSNE plots showing subtypes for WT and NDUFS4 KO separately. **G.** Proportion of subtypes of NSPCs in WT and NDUFS4 KO.

Higher resolution of cell types within the neurogenic lineage was achieved by subclustering NSCs, TAPs, and NBs; resulting in nine neurogenic clusters^21,22^, six of which represent different stem/progenitor states along the neurogenic lineage, such as quiescent vs active NSCs and early vs late NBs **(Figure 4C)**. Subtypes were identified based on expression of (i) known genes (*Aqp4*, *Mki67*, and *Tubb3*) as well as (ii) genes associated with lipid biosynthesis, glycolysis, cell cycle, ribosome, mitosis, and neuronal differentiation **(Figure 4D-E)**. Further proportion analysis within the subclusters revealed a significant reduction in the percentage (33.88% vs 1.75%) of cells identified as early NBs paired with an increase in late NBs (2.40% vs 31.99%) in NDUFS4 KO compared with controls **(Figure 4F-G)**. These data suggest a maturational bias in KO mice whereby late NBs represent most of the NB cell pool and agree with our decline seen in NSPCs undergoing the early lineage progression into NBs (SOX2^+^ DCX^+^) – presumably early NBs **(Figure 2)**.

To further validate our findings suggesting proliferative impairments within the neuronal lineage, we compared the gene expression data sets of NSCs and TAPs for the cumulative fraction of genes associated with the G2/M border and M phases of the cell cycle. NDUFS4 KO NSCs and TAPs exhibited significantly lower cumulative expression of genes associated with these cell cycle phases **(Supplementary Fig. 3).** Paired with our previous evidence of hampered NSPC proliferation *in vivo* and *in vitro*, our findings suggest that NDUSF4 is a critical regulator of neurogenesis during early postnatal development that may be required for the production and maturation of key cell types within the neural lineage occupying the SVZ.

### Single cell transcriptomic profiling demonstrates several dysregulated pathways along the neuronal lineage in NDUFS4 KO mice

To determine which signaling pathways may underlie our observed defects in neurogenesis, we analyzed differential gene expression profiles across the three main subclasses of cells impacted along the neuronal lineage in NDUFS4 KOs: NSCs, TAPs, NBs. We identified over 825 differentially expressed genes for NSCs and 613 genes for TAPs between genotypes (**Figure 5A-D**, **Table S1 & S2).** These differentially expressed genes (DEGs) collectively rank neurogenesis as one of the top two most affected biological processes in NDUFS4 KO NSCs and TAPs, as shown by gene ontology (GO) analysis **(Figure 5B, D)**. Ingenuity pathway analysis (IPA) also revealed dysregulated genes in the KO NSCs associated with neurodevelopmental disorders; these genes have overlapping functions in neural cell proliferation, development, and differentiation **(Figure 5E-F)**.

**Figure 5.**
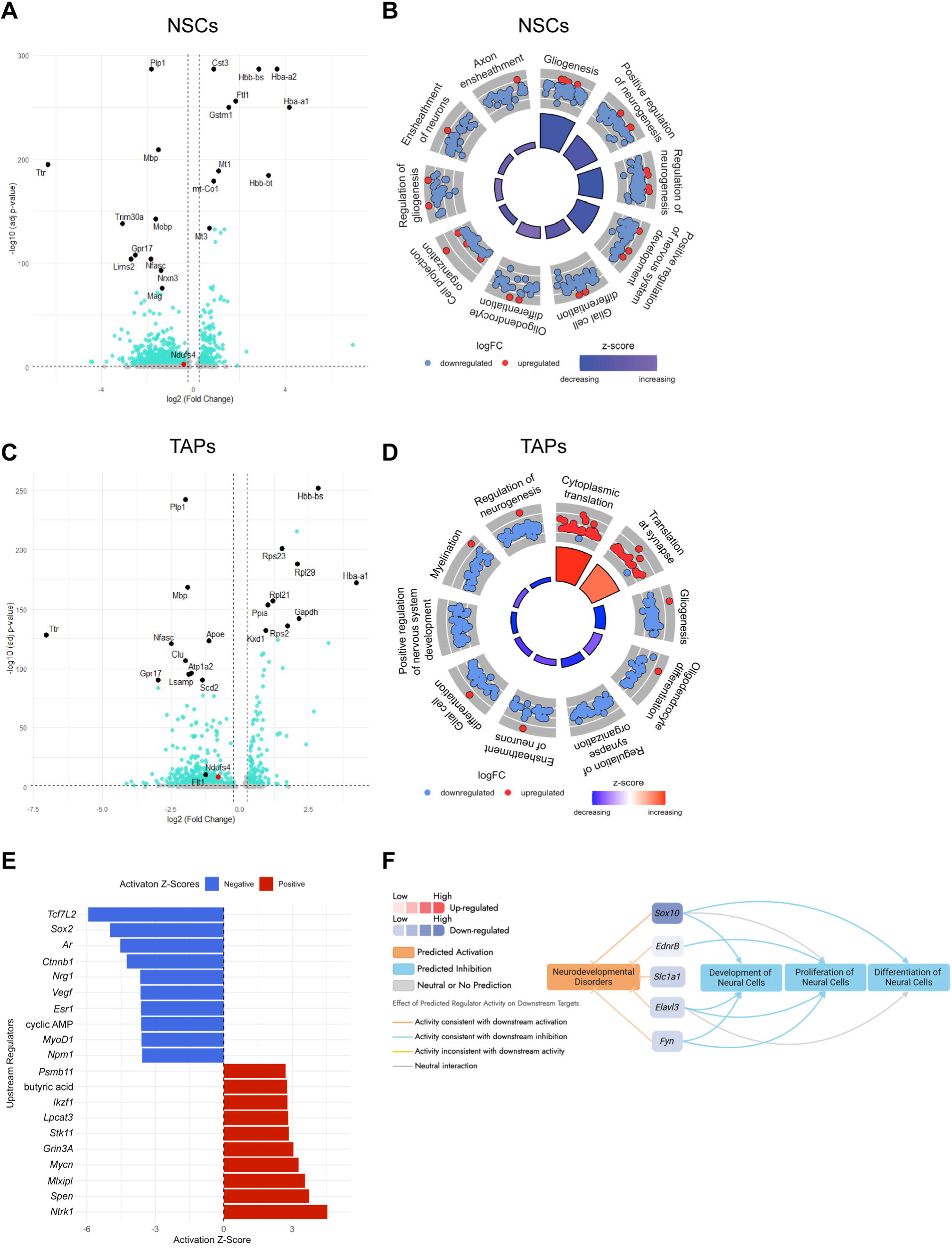
Differentially expressed genes and dysregulated biological pathways in NDUFS4 KO NSCs and TAPs. Volcano plot showing fold change of differentially expressed genes in NDUFS4 KO **A.** NSCs and **C.** TAPs. Top 10 up and downregulated genes are labelled (Black circles) and *Ndufs4* gene is shown in red circle. The thresholds were set at log2(Fold Change) ≥ +0.25 and ≤ -0.25, and – log10 (adj p-value) < 1.2991. GO circle plot showing top dysregulated biological processes and regulation of associated genes in NDUFS4 KO **B.** NSCs and **D.** TAPs. **E.** Top 10 up and downregulated upstream regulators in KO NSCs from ingenuity pathway analysis. **F.** Dysregulated genes in KO NSCs are associated with predicted neurodevelopmental disorders that overlap with neural cell proliferation, development, and differentiation.

Focusing on the neuronal lineage, we found a diverse range of pathways altered at different stages throughout immature neuronal production. For example, we observed upregulation of *Mt1* and *Mt3* in NSCs (**Figure 5A**), which are genes previously reported to be highly expressed in radial glia precursors transitioning into quiescence during late embryonic stages^23^. An upregulation in these genes suggests a possible premature shift toward quiescence in NDUFS4 KO NSCs. We also identified dysregulated upstream regulators of NSC survival (*Tcf7l2* and *Ctnnb1*, involved in Wnt signaling), stem cell self-replication (*Sox2*), proliferation (*Ar*, *Esr1*, *Npm1*, and *Mycn*), and quiescence (*Stk11*)^24-26^ either as a DEG or predicted from IPA **(Figure 5, Table S1)**. Interestingly, DEGs in NSCs also revealed an increase in some subunits of Complex II-IV, which may indicate a compensatory mechanism^27,28^ in the absence of complex I activity, where NDUFS4 is significantly decreased **(Figure 5A, C; Supplementary Fig. 4)**. Additionally, we observed downregulation of *Ttr* in both NSCs and TAPs, which has been implicated in neural stem cell fate decisions^29^. The KO NBs displayed increased expression of 388 genes, mostly associated with increased translation, which may account for the observed push towards late stage neuroblasts^30^ (**Supplementary Fig. 5, Table S3**).

Owing to the diverse classes and subclasses of cells identified, we next wanted to evaluate potential alterations in intercellular interactions. A disruption in cell-to-cell communication was inferred between ligand and receptors upon further bioinformatic analyses between all identified cell types. NDUFS4 KO TAPs and other cell types such as astrocytes had reduced interactions, resulting in changes in the overall information flow between cell types **(Supplementary Fig. 6)**. For example, we observed an upregulation of signaling associated with neural cell adhesion molecules (NCAMs) **(Supplementary Fig. 6)**, which has been implicated in decreased progenitor proliferation in the neurogenic stem cell niche^31^. These analyses help contextualize the cellular milieu within the SVZ in NDUFS4 KO animals whereby loss of complex I activity in mitochondria has not been restricted to a single cell type; therefore, it is also important to consider potential confounds from extrinsic signaling mechanisms. Together, these findings suggest that the postnatal neurogenic niche is changed in composition, altering the genetic profile of the stem cell pools thus resulting in a completely different neurodevelopmental signaling signature in LS than in neurotypical development. We next focused our attention on the second most affected biological process in NDUFS4 KO NSCs: gliogenesis.

### Decreased oligodendrogenesis in NDUFS4 KO mice

Because (i) neurons and oligodendrocytes (OLs) are both derived from NSPCs, (ii) our bioinformatic assessments indicated a significant downregulation in gliosis within the NSC and TAP populations, and (iii) we found a significant reduction in NSPC numbers throughout the SVZ **(Figures 2, 5, Supplementary Fig. 1)**, we reasoned that oligodendrogenesis and functional maturation into myelin producing OLs would be effectively downregulated in the absence of Complex I activity. Recently, remodeling of mitochondrial content, dynamics, and organization has been reported during oligodendrocyte development *in vivo*^32^. Considering that OLs increase in surface area by up to 6,500-fold during maturation/myelination^33^, additional insight into the role mitochondria play during these processes are warranted.

We first performed immunohistochemistry to assess the total number of OLs throughout the entire OL lineage within the SVZ. Quantification of OLIG2^+^ cells exhibited no significant difference in the density of OLs between WT and NDUFS4 KOs **(Supplementary Fig. 7A, C)**. We next assessed the oligodendrocyte progenitor (OPCs) population and found no differences in PDGFRα^+^ OPCs **(Supplementary Fig. 7B-C)**. While OPCs are generated within the SVZ, they must migrate to target axonal tracts to commence maturation and functional myelination; in addition, OPCs are capable of self-renewing following emigration from their points of origin^34^. In considering the dynamic cell classes and transcriptional profiles generated when clustering for neuronal lineage **(Figure 4C)**, we reasoned that there may be an imbalance in specific subclasses of OLs that wouldn’t be accounted for by total cell population analyses. Therefore, we utilized a combination of birth dating transcription factors to assess the embryonic origins and predictive cell fates of stem/progenitor cells within the SVZ.

While oligodendrogenesis commences embryonically (∼E12.5), myelination of the mouse brain occurs postnatally and is critical for proper motor and cognitive functions^35^. There are three known waves of oligodendrocytes derived from distinct NSPC populations at different stages of perinatal brain development; in addition, these waves immigrate from different regional sources including the lateral ganglionic eminence (LGE), medial ganglionic eminence (MGE), and the postnatal cortex^35^. Importantly, NSCs in the postnatal brain are in part derived from embryonic LGE/MGE, maintain their spatial organization (LGE, lateral wall, MGE, ventral tip) (**Figure 6A**), and express some of the same markers^36,37^. Therefore, we performed IHC targeting the transcription factors MASH1 (LGE and MGE) and Nkx2.1 (MGE), at P14, a time in development when the corpus callosum is heavily populated with MGE-derived (∼50%) cells of the OL lineage.

**Figure 6.**
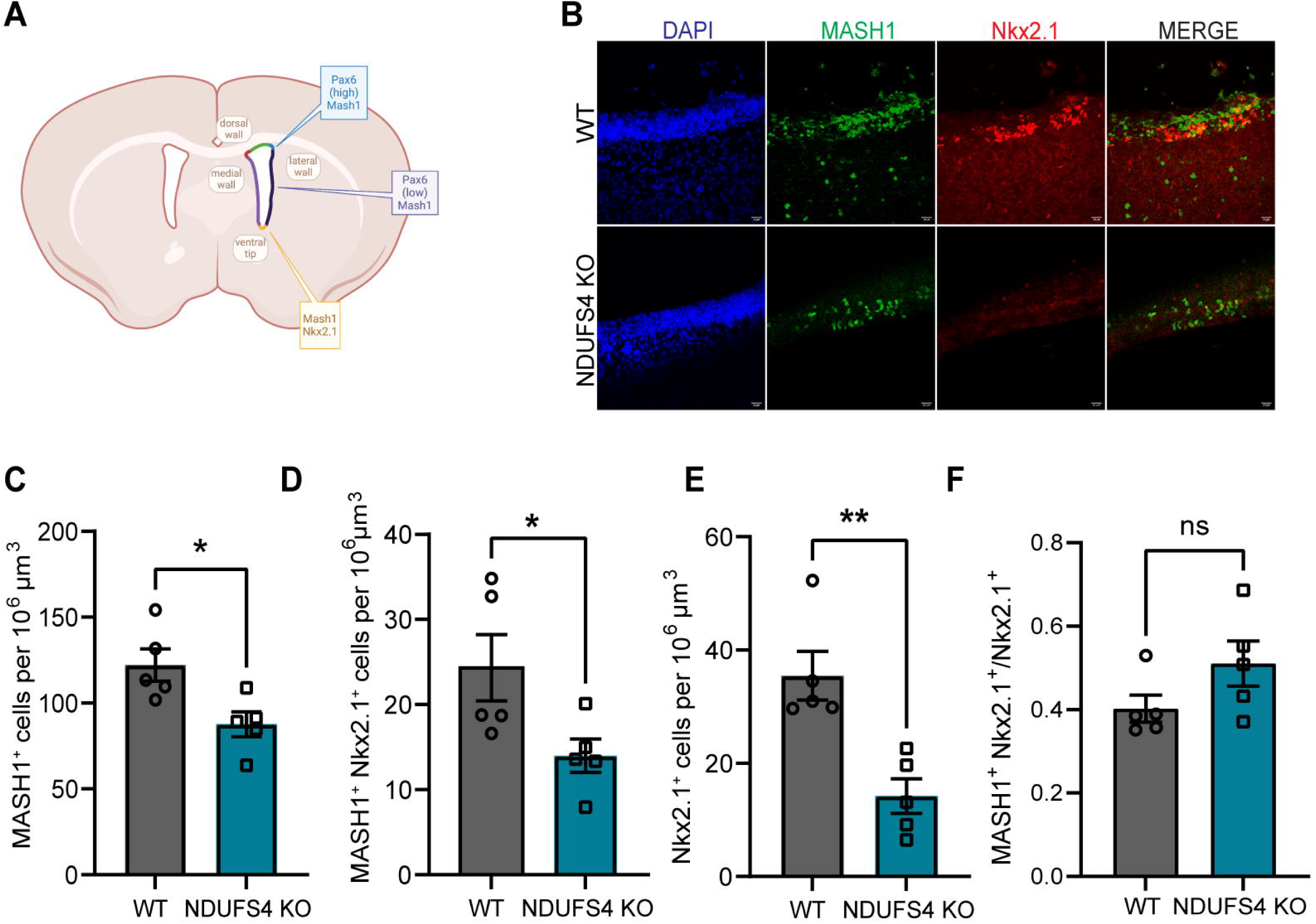
NSCs deriving from the embryonic lateral and medial ganglionic eminences were reduced in the postnatal SVZ of NDUFS4 KO mice. **A.** Schematic illustrating the expression patterns of MASH1, Pax6, and NKX2.1 in the lateral, ventral, and dorsal walls of the SVZ. Made with BioRender. **B.** Representative confocal images of immunohistochemical staining for MASH1 (green), Nkx2.1 (red), and DAPI (blue) in the lateral SVZ of WT and NDUFS4 KO mice at P14. Scale bar = 20 µm. **C–E.** Quantification of cell densities in the SVZ at P14: **C.** MASH1 cells, **D.** double positive Nkx2.1 MASH1 cells, **E.** Nkx2.1 cells and **F.** Ratio of double positive Nkx2.1 MASH1 cells to Nkx2.1^+^ cells. Unpaired t-test was performed; *p<0.05, **p<0.01. Bar plot represents mean and standard error of mean. Each dot represents one animal. n=5 per group.

We found a significant reduction in Nkx2.1^+^, MASH1^+^, and MASH1^+^ Nkx2.1^+^ cells, indicating that NSCs derived specifically from the LGE (MASH1^+^ Nkx2.1^+^) and MGE (Nkx2.1^+^) were affected in the NDUFS4 KO SVZ **(Figure 6B-E)**. We found no significant differences in the ratio of MASH1^+^ Nkx2.1^+^ to Nkx2.1^+^ (40.23% vs 50.01%), suggesting that these populations were seeded in the SVZ in a similar manner from each embryonic source assessed (e.g. - LGE, MGE), with equivalent proportions of OL committed cells within the NSC pools **(Figure 6F)**. These findings are within anatomically equivalent planes of the LGE and MGE - the predecessor tissues to the SVZ - around a postnatal age in which the source of OLs populating the corpus callosum (CC) – the nearest axonal tract to be myelinated – are shifting from primarily LGE-derived to MGE- and cortically-derived^35^. MASH1 is an essential promotor of oligodendrogenesis and subsequent myelination^38^, and MASH1^+^ progenitors contribute to both neurons and oligodendrocytes postnatally^39^; therefore, a loss in the number of MASH1^+^ cells may have an impact on both oligodendrogenesis and neurogenesis, defining a cell type that lies at the intersection of the mechanism by which Complex I deficiency results in brain maturational delays evident prior to the onset of the neurodegenerative signatures in LS.

### Complex I deficiency results in disrupted myelination of corpus callosum

Because MGE-derived OPCs migrate to and myelinate the corpus callosum, and we found a significant reduction in PLP1 gene expression - the most abundant myelin protein in the brain - in our single cell seq data **(Figure 5A)**, we next assessed the CC. Corpus callosum formation begins prenatally whereby the initial axons cross the brain midline and continues through early postnatal days, with migrating OPCs from the dorsal wall populating this axonal tract and generating mature myelinating oligodendrocytes^40^. We reasoned that an impairment in either of these processes - neurons sending axons through the CC or loss of myelin - would result in an overt anatomical phenotype.

Considering our findings of reduced brain hemispheric width in KO animals (**Figure 1C-D**), we measured landmark neuroanatomical structures on the coronal planes corresponding with the Bregma areas where we found the most significant differences (+1.53 and +1.23) **(Supplementary Fig. 1)**. We found no differences in cortical mantle thickness, striatal width, or SVZ volume **(Figure 7A-C)**. When assessing the CC, we found a significant reduction in thickness in NDUFS4 KO compared to WT **(Figure 7D, E)**. We next performed immunostains for PLP1 to visualize myelin **(Figure 7F)**. Quantification of mean fluorescence intensity **(Figure 7G**), percentage area (**Figure 7H-I**) – to account for differences in CC size – covered, and mean area coverage **(Figure 7J)** revealed a significant reduction in PLP expression and quantity, indicating a disruption in myelination **(Figure 7G-J)**. In addition, probing for myelin basic protein (MBP), another structural protein critical for myelination, revealed similar expression patterns and significant reduction **(Figure 7K-L)**. Together, these findings indicate that Complex I activity is critical for OL production and myelination within the postnatal SVZ and CC. While agenesis of the CC has been documented in a few LS cases, the areas of the brain most assessed and affected are deep brain structures seeded with neurons early in embryonic development^41^. While myelination is strictly a postnatal phenomenon in the mouse brain, OL lineage progression follows a series of steps akin to neurons which vary in energetic demand at each stage. Therefore, diffuse white matter injury may represent a largely overlooked phenotype in monitoring human neonatal brain development and a defect that precedes the pediatric neurodegenerative manifestations most ascribed to motor and cognitive impairments in LS.

**Figure 7.**
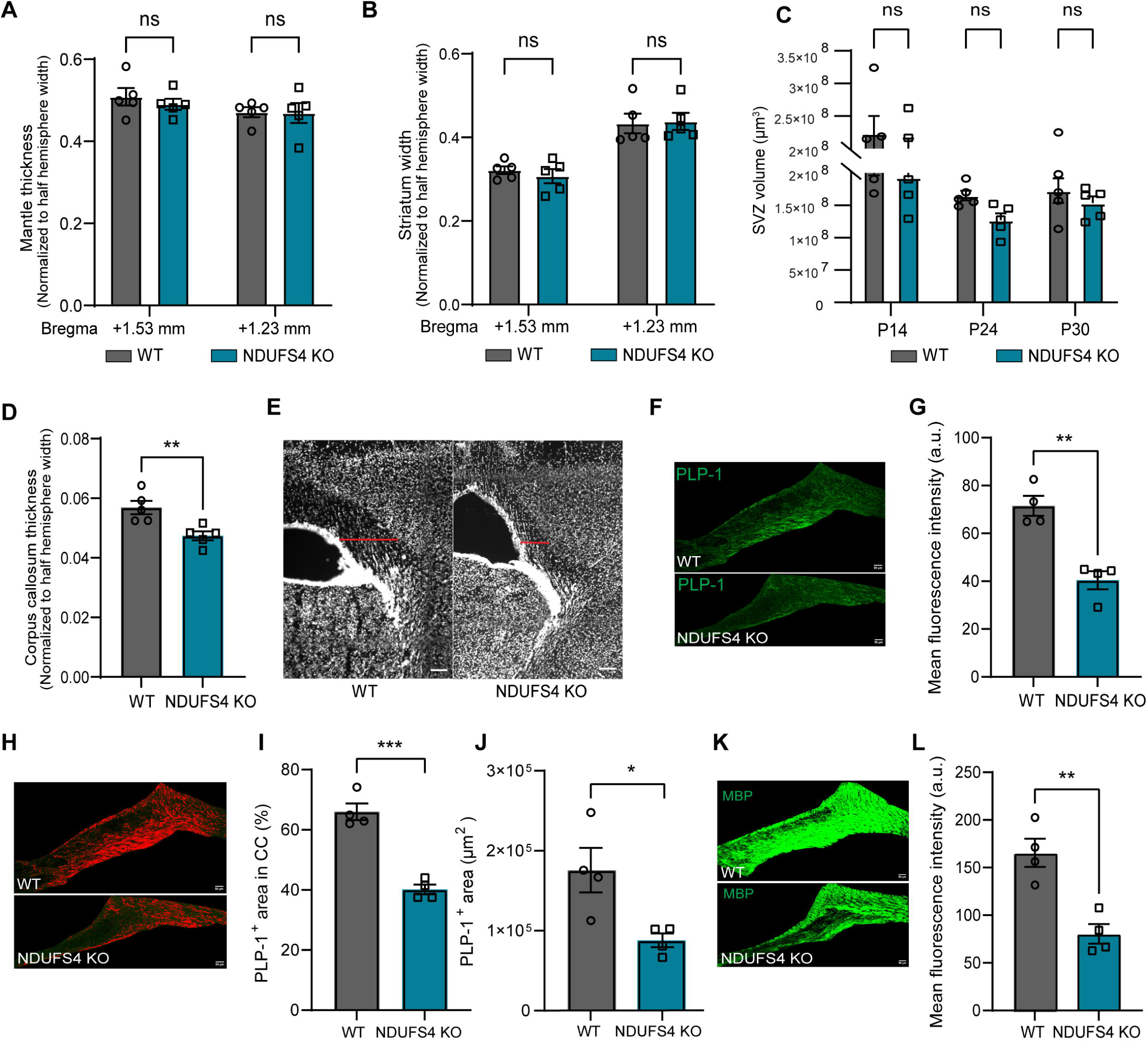
Reduced Myelination in NDUFS4 KO corpus callosum. **A-D.** Measurements of neuroanatomical structures from coronal sections at +1.53 mm and +1.23 mm from bregma: **A**. Mantle thickness, **B.** Striatum width. **C.** Estimated SVZ volume measured across five coronal sections per animal. Two-way ANOVA was performed; ns= not significant. **D.** Corpus callosum thickness measured from coronal sections at +1.23 mm from bregma. **E.** Representative confocal images of the corpus callosum from WT and NDUFS4 KO mice at P14. Scale bar= 100 µm. Red dotted line represents measurement taken for this image. **F.** Representative confocal images of immunohistochemical staining for PLP-1 in corpus callosum. Scale bar= 50 µm. **G.** Quantification of mean fluorescence intensity of PLP-1. **H.** Threshold optimized confocal images of WT and NDUFS4 KO corpus callosum presenting area coverage by PLP-1 fluorescence. Scale bar= 50 µm. **I.** Percentage of area covered by PLP-1 fluorescence in corpus callosum. **J.** Mean area coverage of PLP-1 in CC. **K**. Representative confocal images of immunohistochemical staining for MBP in corpus callosum. Scale bar= 50 µm. **L.** Quantification of mean fluorescence intensity of MBP in corpus callosum. Unpaired t-test was performed; *p<0.05, **p<0.01, and ***P<0.001. Bar plot represents mean and standard error of mean.

## Discussion

Using the preclinical mouse model of LS, we investigated the neurodevelopmental underpinnings of the disease considering that new reports indicate patients do not solely display degenerative regressive symptomology. Our work demonstrates that prior to appearance of neurodegenerative phenotypes, neural stem and progenitor function and differentiation is abnormally perturbed postnatally and the temporal timing of typical neurodevelopmental events are delayed during LS. Our data indicates that the disruption of NDUFS4 disrupts cell type composition of neurogenic niches as well as the transcriptomic signatures of the NSPCs altering the phenotypes of these cells.

Previous reports have indicated the sensitivity of stem cell biology to mitochondrial dysfunction in models of detrimental mitochondrial DNA damage or other types of unphysiologically disruptions of mitochondrial function. In models with double strand break damage to mitochondrial DNA (mtDNA), highly proliferative stem cell populations appeared most affected^42,43^. Neural and hemopoietic stem cell defects were described in the Mutator mice, where a proofreading defective polymerase γ causes a high level of mtDNA mutations^44,45^. The loss of apoptosis-inducing factor (AIF), a mitochondrial inner membrane flavoprotein, severely disrupts complex I cause embryonic lethality^46^. Conditional ablation under the control of NSPC promoters of AIF or mitochondrial fission or fusion genes, such as *Mfn1/Mfn2* and *Drp1*, halt the differentiation process and ability for self-renewal^47^. However, considering that these studies have now provided the basis for research into the links between stem cell and mitochondrial function, different models that better replicate disease conditions where mitochondrial function is perturbed are required to understand the pathophysiological impact to stem cell biology.

Neural stem cells are more reliant on glycolysis and ramp up oxidative metabolism during differentiation^51,52^. However, recent evidence demonstrates that mitochondrial dysfunction results in abnormal neurogenesis. Defects in neurogenesis contribute to cognitive and social defects associated with neurodevelopmental disorders such as Williams and DiGeorge/22q11 deletion syndrome^53,54^. In another neurodevelopment disorder schizophrenia, the biggest genetic risk factor, 3q29Del, overwhelmingly points to alterations in mitochondrial DNA transcripts and nuclear genes that control OXPHOS in induced pluripotent stem cell (iPSCs) organoid and mouse models^55^. These data and our own indicate that mitochondrial dysfunction affects not only the differentiation process and post-mitotic neuronal function, but properties of the NSPCs itself^48^. Altering the ability to generate new neurons development or even adulthood in the hippocampus and SVZ, or passing on damaged mitochondria to either stem, progenitor or differentiated progeny, could have lifelong negative outcomes and consequences.

Neurodegenerative disease-causing genetic mutations leading earlier neurodevelopmental perturbations are overlooked, but most likely not surprising. The human SVZ remains neurogenic, with an abundance of newborn interneurons migrating through the SVZ to the prefrontal cortex throughout the first 1.5 years of life^49^. For example, the neurodegenerative disease Huntington’s disease (HD) is due to polyglutamine expansion in the *HTT* gene manifesting in one’s middle to late adulthood affecting the striatum and basal ganglia^50^. However, high repeat expansions lead to juvenile onset HD displaying different symptomology and neuroanatomic regions affected, than adult HD^51^. These high number of repeats modeled in iPSCs, or human embryonic stem cells (hESCs) display abnormal neuronal and cortical differentiation and disrupted neurogenesis^52,53^. This is also evident in an Alzheimer’s disease (AD) mouse model that harbors familial mutations that display reductions neurogenesis and abnormal neurodevelopment in the subgranular zone of the hippocampus as early as P7^54^. Granted, most forms of AD are sporadic, so neurodevelopmental defects may not always precede this disease, but the investigation into the neurodevelopmental underpinnings of neurodegenerative disease phenotypes and variation in symptomology is warranted.

While lesions within the gray matter are evident in LS patients and recapitulated in mouse models, less attention has been given to the white matter essential for neurotypical motor and cognitive development. With consideration of how neuronal and oligodendroglial developmental programs unfold in sequence in mice, our collective findings - reduced Mash1^+^ OLs in the SVZ, no changes in SVZ volume, and no changes in total OL population within the SVZ - indicate a disruption (i) in local oligogenesis within the CC, (ii) stall in lineage progression, (iii) inability to generate myelin, (iv) loss of myelin initiating cues from unhealthy/dysmature axons or a combination of cellular stress points. Further conditional genetic studies paired with axonal tracers will greatly aid in deciphering the full impact of the eclectic changes in transcriptomic profiles identified. While it is known that each of the 3 competitive waves of OPC development can functionally replace one another^35^, which may explain why we only see differences in a proportion of OPCs rather than whole population (PDGFRα), our PLP findings suggest that this phenotype is more representative of a failure to mature or myelinate the CC subsequent from migration from SVZ.

Dysmyelination of the CC may be dampened in humans with LS compared to this mouse model as humans born with a mixed profile of OLs including progenitors, as myelination commences during the 2^nd^ trimester^55,56^; therefore, the stress imposed by their *ex utero* environment may manifest in delays in myelination rather than complete lack of corpus callosum formation. Recent evidence demonstrates that loss of the *Nfia1* gene results in agenesis of the corpus callosum^57^ and our transcriptomics demonstrate a significant reduction in this gene within the TAPs suggesting that this progenitor pool, capable of producing oligodendrocytes and neurons, may react to complex I deficiency differently before and after birth. The CC is a relatively small structure to quantify by standard clinical MRI assessments in the young and, unlike more noticeable cystic white matter lesions, may require imaging modalities such as diffusion tensor imaging (DTI) to accurately assess diffuse changes in myelination/integrity. Such prospective clinical studies in LS patients in parallel with well-designed/characterized genetic murine models of mitochondrial dysfunction in the young will be invaluable in developing preventative and predictive treatment strategies to improve psychomotor outcomes.

### Limitations of the study

We found that postnatal neurodevelopmental processes were disrupted in a preclinical LS mouse model. More thorough examination embryonically and at the other postnatal neurogenic niche in rodents, like the hippocampus, is warranted. Another study knocking out NDUFS2 under the control of the overexpressed transgenic human *GFAP* promoter described neurogenic postnatal defects, but the model produced a severe impact embryonically starting at day 13.5^58,59^. It is also unclear if the severity of the defects described in this study extend to LS caused by other mutations that are not Complex I specific. However, we highly find that this is likely due to recent work describing NSPC defects in an iPSC organoid model harboring *SURF1* mutations^60^.

## Abbreviations

AD: Alzheimer’s disease
CC: corpus callosum
DCX: doublecortin
IHC: immunohistochemistry
GO: Gene Ontology
hESCs: human embryonic stem cells
HD: Huntington’s disease
KO: knockout
iPSCs: induced pluripotent stem cells
LGE: lateral ganglionic eminence
LRT: likelihood ratio test
LS: Leigh syndrome
MGE: medial ganglionic eminence
NBs: neuroblasts
NSCs: neural stem cells
NSPCs: neural stem progenitor cells
OL: oligodendrocyte
OXPHOS: oxidative phosphorylation, postnatal day (P#)
scRNA: single cell RNA
SVZ: subventricular zone
TAPs: transit amplifying cells
t-SNE: t-distributed Stochastic Neighbor Embedding
UMAP: Uniform Manifold Approximation and Projection
WT: wildtype

## Author Contributions

SRB planned and performed experiments, analyzed data, and wrote and revised the manuscript. PLT, BAL, JPM, SB, ND, SNH, YS, and CK performed experiments. PDM and AMP conceived project, planned experiments, analyzed data, and wrote and revised the manuscript. All authors read and approved the manuscript.

## Acknowledgments

We thank the National Institutes of Health R01ES035013 (PDM) and R35GM142368 (AMP) for supporting this work.

## Declaration of Interests

The authors declare no competing interests.

## STAR Methods

### Lead Contacts and Materials Availability

Further information and requests for resources and reagents should be directed to and will be fulfilled by Lead Contacts, Paul D. Morton, Ph.D. and Alicia M. Pickrell, Ph.D.

### Animals

Wild-type (WT) and heterozygous (Het) mice for the *Ndufs4* gene (B6.129S4-*Ndufs4^tm1.1Rpa^/J*) were obtained from Jackson Laboratory. Mice were housed under standard conditions, maintained on a 12-hour light/dark cycle, with ad libitum access to food and water. To generate *Ndufs4* knockout (KO) animals, heterozygous *Ndufs4* mice were crossed. For immunohistochemical analysis, mixed sex was used. Single cell sequencing samples were collected from the following sexes: I) WT1 was derived from one male and one female WT mouse, II) WT2 was derived from a single female WT mouse, III) NKO1 and NKO2 were derived from two different female NDUFS4 KO mice. All experimental procedures were conducted in accordance with the NIH Guide for the Care and Use of Laboratory Animals and were approved by the Virginia Tech Institutional Animal Care and Use Committee.

### Immunohistochemistry

To prepare brain sections for IHC, mice were anesthetized with a ketamine/xylazine cocktail and perfused with cold 1X PBS containing 1,000 IU/mL heparin. The brains were harvested and incubated in 4% paraformaldehyde (PFA) overnight at 4°C. After fixation, the brains were incubated with 30% sucrose in PBS for 24 hours until the brain sunk and then frozen in OCT media (Tissue-Tek) containing 30% sucrose. Frozen brains were sectioned and mounted at a thickness of 30 µm and stored at –80°C until further use. For antigen retrieval, sections were incubated in 95°C sodium citrate buffer (pH 6) for 5 minutes. Tissue sections were permeabilized with 0.4% Triton X for 10 minutes followed by incubation in blocking buffer (5% Bovine serum albumin + 2.5% Normal goat serum) for 1 hour. Tissue sections were incubated with primary antibodies at the following dilutions: rabbit anti-Sox2 (1:250; MilliporeSigma #AB5603), guinea pig anti-DCX (1:350; MilliporeSigma #AB2253MI), mouse anti-MASH1 (1:200; BD Biosciences #556604), rabbit anti-Nkx2.1 (1:300; Fisher Scientific #07601MI), mouse anti-Olig2 (1:250, EMD millipore #AB9610), rat anti-PDGFRα (1:200; BD Pharmigen #558774), mouse anti-Ki67 (1:250; BD Pharmigen #550609), rabbit anti-Myelin PLP (1:500, abcam #ab28486), and rabbit anti-Myelin Basic Protein (1:50, Cell signaling Technology #78896). All secondary antibodies were used at a 1:200 dilution. A standard tyramide signal amplification (TSA) protocol (ThermoFisher Scientific) was employed as per manufacturer’s instruction whenever primary antibodies from the same host species were used. Stained tissues were mounted using DAPI Fluoromount-G (SouthernBiotech #0100-20).

### Imaging

Following immunostaining, high-resolution Z-stack images (1 µm step size) of the dorsolateral, lateral and dorsal (for PDGFRα and Olig2 staining) walls of the subventricular zones (SVZs) were captured using a Nikon C2 confocal laser scanning microscope (Nikon Instruments, Melville, NY) at 40X magnification. Reference maps at 10X magnification were used to guide the acquisition of the 40X images. Neurospheres were imaged under a brightfield filter using a 10X objective.

The Nikon C2 is outfitted with brightfield and phase contrast with a Kinetix CMOS camera and fluorescent channels DAPI, GFP, Texas Red, and Far Red. The Nikon LUN4 has a four-line solid-state laser system with Perfect Focus and DU3 High Sensitivity Detector System. The CFI60 Apochromat Lambda S 40× water immersion objective lens, N.A. 1.25, W.D. 0.16–0.2 mm, F.O.V. 22 mm, DIC, correction collar 0.15–0.19 mm, spring-loaded was used for confocal image acquisition. The Kinetix CMOS camera and 10X objective was used for bright field image acquisition. Nikon NIS-Element Package and ImageJ were used for image analysis.

### Cell quantification

Target cells in the dorsolateral, lateral, and dorsal regions of the subventricular zone (SVZ) were exhaustively quantified from five coronal sections per animal. Sections were spaced 300 µm apart, covering the entire SVZ. All cell counting was performed using ImageJ software.

### Statistical analysis for IHC

Student’s t-test was used for comparisons between WT and KO groups at a single time point. For comparisons across multiple time points, a two-way ANOVA was performed. A p value < 0.05 was considered statistically significant.

### SVZ tissue collection and single-cell dissociation

For single-cell experiments, P14 mice were anesthetized and hand perfused with ice cold PBS. Brains were promptly harvested and placed in ice-cold HBSS without Ca^2+^ and Mg^2+^. The lateral wall of the subventricular zone (SVZ) was dissected following the protocol^61^.

Tissue was dissociated using the Neural Tissue Dissociation Kit (P) (Miltenyi #130-092-628), as described by^68^. Briefly, the dissected tissue was minced on ice using a #10 scalpel blade and transferred to enzyme mix 1. This mix consisted of buffer X, 70 μM beta-mercaptoethanol, 0.04% BSA, and enzyme P (added immediately before use). The tissue was incubated at 37°C for 15 minutes with intermittent pipetting to aid in dissociation. Afterward, enzyme mix 2 (buffer Y and enzyme A) was added, and the tissue was incubated for an additional 10 minutes with further pipetting steps. Dissociation was stopped by adding HHB solution (HBSS containing calcium, magnesium, 10 mM HEPES, and 0.01% BSA), and the cell suspension was filtered through a 70 μm strainer.

The filtered cell suspension was centrifuged at 500×*g* for 10 minutes at 10°C. Cells were resuspended in HHB and purified using a three-layer Percoll gradient (19%, 15%, and 11% Percoll in HBSS/HEPES). Following centrifugation at 430×*g* for 3 minutes at 4°C, the top debris layer was discarded, and the remaining cells were pelleted, washed, and filtered through a 30 μm strainer. Cell counts were performed using trypan blue, and the concentration was adjusted to 5000 live cells/μL in cell suspension buffer provided by PIPseq (Fluent Biosciences) for subsequent cell lysis and library preparation as per manufacture’s instruction.

### Cell preparation

Single-cell suspensions were prepared and diluted to 5,000 cells/µL in Cell Suspension Buffer (Fluent Biosciences, FB0002440), with a total of 40,000 cells per reaction. Each suspension was supplemented with 40U of RNase inhibitor (MilliporeSigma #3335399001).

### Capture and lysis

T20 PIPs (FB0003914) were thawed, briefly centrifuged, and placed on ice. Each T20 PIPs (FB003914) tube received 8 µL of the cell suspension, 40U of RNase inhibitor, and 1 mL of Partitioning Reagent (FB0003123). After mixing and vortexing, the excess Partitioning Reagent was removed, leaving the PIPs+cells. Chemical Lysis Buffer 3 (CLB3, FB0003910) was added to the PIP tubes and the samples were incubated in preheated PIPseq™ Dry bath (FBS-SCR-PDB, 25°C for 15 mins- 37°C for 45 mins- 25°C for 10 mins), to complete lysis.

### mRNA isolation

Following lysis, PIPs were processed with Breaking Buffer (FB0003128) and De-Partitioning Reagent (FB0002516). PIPs were washed three times with chilled 1X Washing Buffer (FB0003139) using centrifugation at 750 x g for 2 minutes. The final PIP volume was normalized to 250 µL based on weight to ensure consistency across samples.

### cDNA synthesis

Reverse transcription was performed with RT Enzyme Mix (FB0002206), RT Additive Mix (FB0002205), and Template Switching Oligo (TSO, FB0003140) in a total reaction volume of 250 µL. Dry bath lid was set to 105°C and the following program was used for cDNA synthesis: 25°C for 30 minutes, 42°C for 90 minutes, and 85°C for 10 minutes.

### cDNA amplification

Whole transcriptome amplification (WTA) was conducted using a master mix comprising 4X PCR Master Mix (FB0004644) and WTA Primer (FB0002006). Amplification reactions were cycled according to protocol, yielding amplified cDNA for downstream library preparation.

### cDNA isolation and QC

Amplified cDNA was isolated from PIPs using magnetic bead cleanup. The purified cDNA was quantified using a Qubit 1X dsDNA High Sensitivity Assay Kit (Thermo Fisher, Q33230), and fragment size distribution was assessed using a BioAnalyzer High Sensitivity DNA Kit (Agilent, 5067-4626).

### Library preparation

Purified cDNA was processed through fragmentation, end-repair, adapter ligation, and indexed PCR using reagents from the PIPseq T20 3’ RNA Kit. Cleanup was performed after each step, with a double-sided size selection following indexing PCR. Libraries were assessed for concentration and quality using Qubit and fragment analysis prior to sequencing. This workflow provided sequencing-ready libraries within two days, suitable for downstream gene expression analysis and single-cell clustering.

### Sequencing

Premade libraries were sent to MedGenome for sequencing on the NovaSeq X+ platform. Libraries were sequenced using 10B lanes with a target of 400 million paired end reads per library and a sequencing depth of 20,000 reads per input cell.

### Data preprocessing

Barcodes identifying individual PIPs/cells were assigned to sequenced cDNA constructs using PIPseeker™ v3.3. PIPseeker first identified barcode combinations and molecular identifiers (MI) for each read before mapping with STAR. After gathering molecule information, reads were assigned to genes by aligning coordinates against the STAR reference, with duplicate reads sharing the same MI collapsed into single counts.

The resulting counts were organized into a sparse matrix, where each entry corresponded to a cell barcode and gene. Barcodes were categorized as either cell-containing PIPs or background PIPs using a barcode rank plot ("knee plot"), which orders barcodes by transcript count. Following cell calling, directories were created for each sensitivity level, containing text files with selected cell barcodes, knee plot images, and filtered count matrices for downstream analysis.

### Initial QC and clustering

To maximize cell inclusion, we used the filtered gene expression matrix generated with the lowest sensitivity threshold. Additional cell filtering criteria were applied based on mitochondrial genome content (<10%) and the number of detected features (>500 and <6000) using Seurat (v1.4). Data normalization, scaling, and variable feature selection were conducted using SCTransform, which effectively mitigates technical noise and accounts for differences in sequencing depth. Principal Component Analysis (PCA) was then performed for dimensionality reduction.

To address batch effects, harmony was applied to the normalized data, ensuring that the batch-corrected data maintained biological variance. The corrected dataset was visualized using Uniform Manifold Approximation and Projection (UMAP) for dimensionality reduction and clustering.

Cell clustering was performed using the FindNeighbors and FindClusters functions, with a resolution parameter set to 0.55. Cluster-enriched genes were identified using the FindAllMarkers function with parameters min.pct = 0.25 and logFC.threshold = log(1.2). Cluster identities were assigned by cross-referencing known cell markers from the literature. Heatmaps displaying the top five enriched genes per cluster were generated using the DoHeatmap function. Additionally, violin plots for known cell markers identified ^62^ were generated for further validation of cluster identities.

### Subclustering and gene set scoring

To identify subclusters within specific cell populations, we isolated neural stem cells (NSCs), transit-amplifying progenitors (TAPs), and neuroblasts from our dataset. Subset data were analyzed using Seurat by rerunning the FindClusters function on principal components, with the resolution parameter set to 0.4. The results were visualized using t-distributed Stochastic Neighbor Embedding (t-SNE).

For cell type characterization and comparison with previously published data, we utilized gene sets from^20,22^. Cells were scored using the AddModuleScore function for genes from the downloaded sets that were expressed in more than 50 UMIs across all cells of interest. The expression patterns of the gene set scores were visualized using violin plots.

### Differential gene expression analysis

To identify dysregulated genes in cells of interest derived from the Ndufs4 KO compared to the WT, we performed differential gene expression (DE) analysis for each cell cluster. The analysis was conducted using the FindAllMarkers function (Wilcoxon rank-sum test) in Seurat and edgeR^63^ for more robust statistical evaluation. Each cell was treated as an individual sample, and both replicates were analyzed independently (WT2 vs. NKO1 and WT1 vs. NKO2). Within each cluster, we compared gene expressions between conditions (KO vs. WT) to identify differentially expressed genes. We report genes that met the following criteria: i) adjusted p-value < 0.05, ii) log2 fold change ≥ +0.25 or ≤ -0.25, and iii) consistent directionality of regulation across both replicates. All non-coding RNAs were removed from the analysis. The results were visualized using volcano plots, where the x-axis represents the log2 fold change, and the y-axis represents the -log10 (adjusted p-value.) Significant genes were highlighted to illustrate key findings in the data.

### Genetic ontology and gene network enrichment analysis

Functional enrichment analysis using DAVID was used to identified altered biological processes. Genes with an adjusted p-value < 0.05 and a log2 fold change ≥ +0.25 or ≤ -0.25 were included in the analysis. Additionally, DAVID was used to retrieve corresponding EnsemblIDs for gene identifiers. For visualization, we utilized the GOplot R package to generate a circle plot, highlighting the top Gene Ontology (GO) terms. This visualization integrates fold-change data with GO enrichment results, providing insights into the direction of regulation (upregulated or downregulated) and the associated biological processes.

### Cluster proportion comparison analysis

The proportions of each cluster and subcluster were compared between groups using the scCODA v0.1.9 Python package (https://sccoda.readthedocs.io/en/latest/compositional_data.html). Ependymal cells were chosen as the reference group for cluster analysis, while for subcluster analysis, the reference cell type was automatically assigned by the software.

### Cell-cell communication analysis

CellChat was used to identify altered signaling pathways in each replicate individually by analyzing the information flow of each pathway^64^. The information flow is calculated as the total communication probability between all pairs of cell groups within the inferred network (https://github.com/sqjin/CellChat).

### Proliferation scoring

Proliferation scoring was performed using the AddModuleScore function^65^, analogous to gene set scoring, but focused on genes associated with different phases of proliferation. For each cell cluster and condition, the cumulative fraction of cells was calculated against the proliferation score. To assess statistical significance, the Kolmogorov-Smirnov test was applied to compare the cumulative distributions between the two conditions for each cluster.

### Ingenuity pathway analysis (IPA)

DEGs from NDUFS4 KO NSCs, determined using the thresholds above, were submitted through Ingenuity Pathway Analysis software (Qiagen) for analyses of upstream regulators and identification of affected canonical pathways. The top 10 inhibited and top 10 activated upstream regulators were used to generate the associated bar plot. Neurodevelopmental disorders were selected from predicted diseases and functions output, and visually reconstructed in BioRender. From affected canonical pathways, oxidative phosphorylation was selected, and expression level of respiratory chain complexes were presented using BioRender.

### Neurosphere isolation and assay

To grow neurospheres, dissociated cells from the lateral wall were harvested and cultured in DMEM/F12 medium (Gibco) supplemented with N2 supplement (Fisher Scientific #17502048), EGF (20 ng/μL, R&D Systems # 236-EG-200), and FGF (20 ng/μL, R&D Systems # 233-FB-025/CF) from P14 animals. Cells were plated in ultra-low adherent 24-well culture plates (Fisher Scientific #NC1882826), and half of the media was replaced every other day during a one-week incubation period. After one-week, primary neurospheres were dissociated using Accutase (STEMCELL Technologies #07920), then replated in ultra-low adherent 6-well plates for secondary neurosphere formation. After three days, neurospheres were transferred onto 2-chambered slides, and brightfield images were captured for analysis. The diameters of 50 neurospheres per replicate (three replicates per genotype) were measured using ImageJ. To compare the distribution of neurosphere diameters between WT and NDUFS4 KO, we performed an ordinal logistic regression analysis with a likelihood ratio test (LRT) in R.

### Data and code availability

Single-cell RNA-seq data have been deposited at GEO at GEO: GSE297078 and are publicly available as of the date of publication. Any other raw data sets generated during this study are currently being used for future studies and to obtain grant funding, but data is available upon request. All original code has been deposited at Zenodo at [10.5281/zenodo.15424859] and is publicly available as of the date of publication.

## Notes

### Competing Interest Statement

The authors have declared no competing interest.

