## Supplementary Figures and Legends for "Impaired Complex I dysregulates neural/glial precursors and corpus callosum development revealing postnatal defects in Leigh Syndrome mice"

Biswas *et al.*

This PDF file includes Figure S1- S7.

Table S1-S3

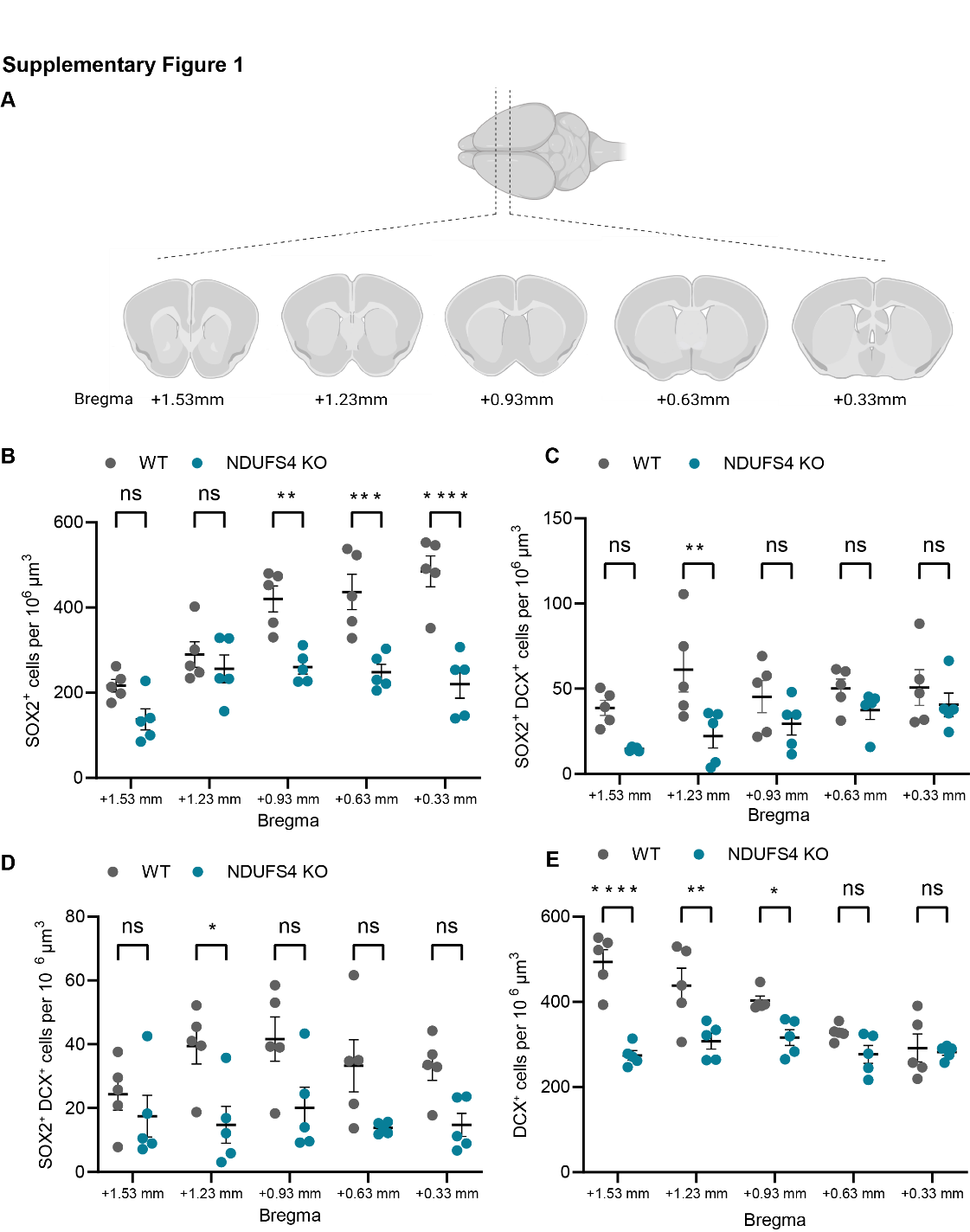

**Supplementary Figure 1. Neural stem and progenitor cell spatial distribution between genotypes.** **A.** Graphical depiction of brain regions analyzed in this study generated with Procreate software. **B** Number of SOX2^+^ cells between WT and NDUFS4 KO mice in the SVZ at P14. **C-D.** Number of SOX^+^ DCX^+^ cells between WT and NDUFS4 KO mice in the SVZ at (C) P24 and (D) P30. **E.** Number of DCX+ cells between WT and NDUFS4 KO mice in the SVZ at P30. Two-way ANONA was performed. *p<0.05, **p<0.01, ***p<0.001, ****p<0.0001, ns= not significant. Each dot represents one animal.

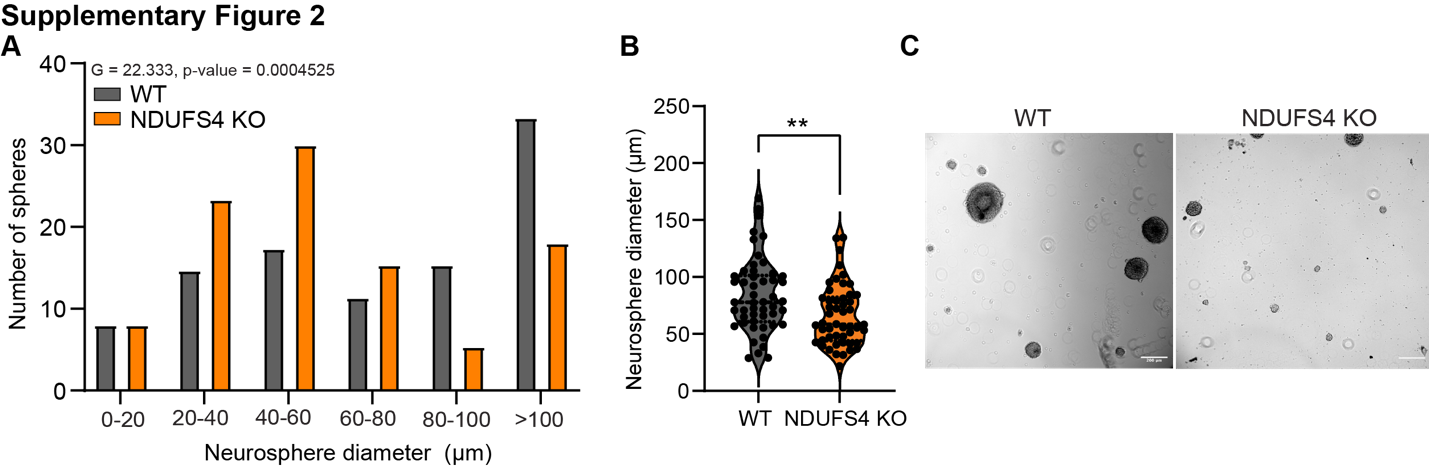

**Supplementary Figure 2. NDUFS4 KO neural stem cells display a reduced ability to proliferate *in vitro*.** **A.** Distribution of neurosphere diameter between WT and NDUFS4 KO animals. Neural stem cells were collected from 3 cultures derived from one animal/genotype. 50 neurospheres were measured per culture. Likelihood ratio test (LRT) was performed to compare the distribution. **B.** Average neurosphere diameter between WT and NDUFS4 KO. **C.** Representative brightfield images for neurospheres. Scale bar = 200 µm. n=150 cells/group. Student’s t-test was performed. **p<0.01. Each dot represents one neurosphere

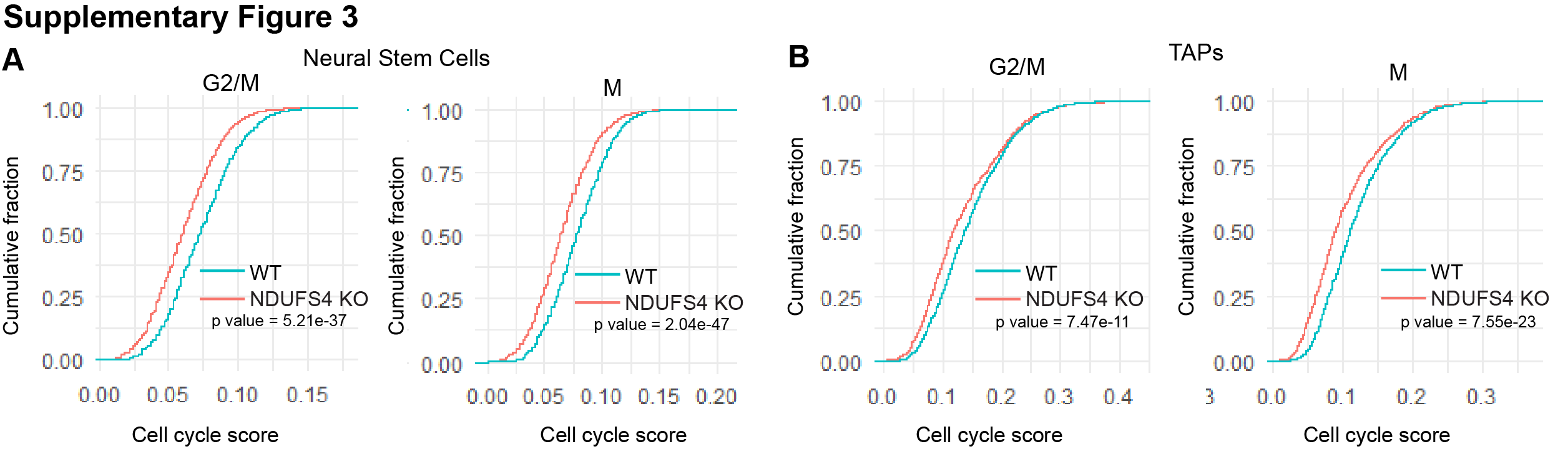

**Supplementary Figure 3.** **Cell cycle scoring from scRNA sequencing data of the SVZ of NDUFS4 KO mice display perturbed proliferation.**

**A-B.** Comparison of cumulative fractions of cell cycle scores for G2/M and M phases between WT and NDUFS4 KO (A) NSCs and (B) TAPs.

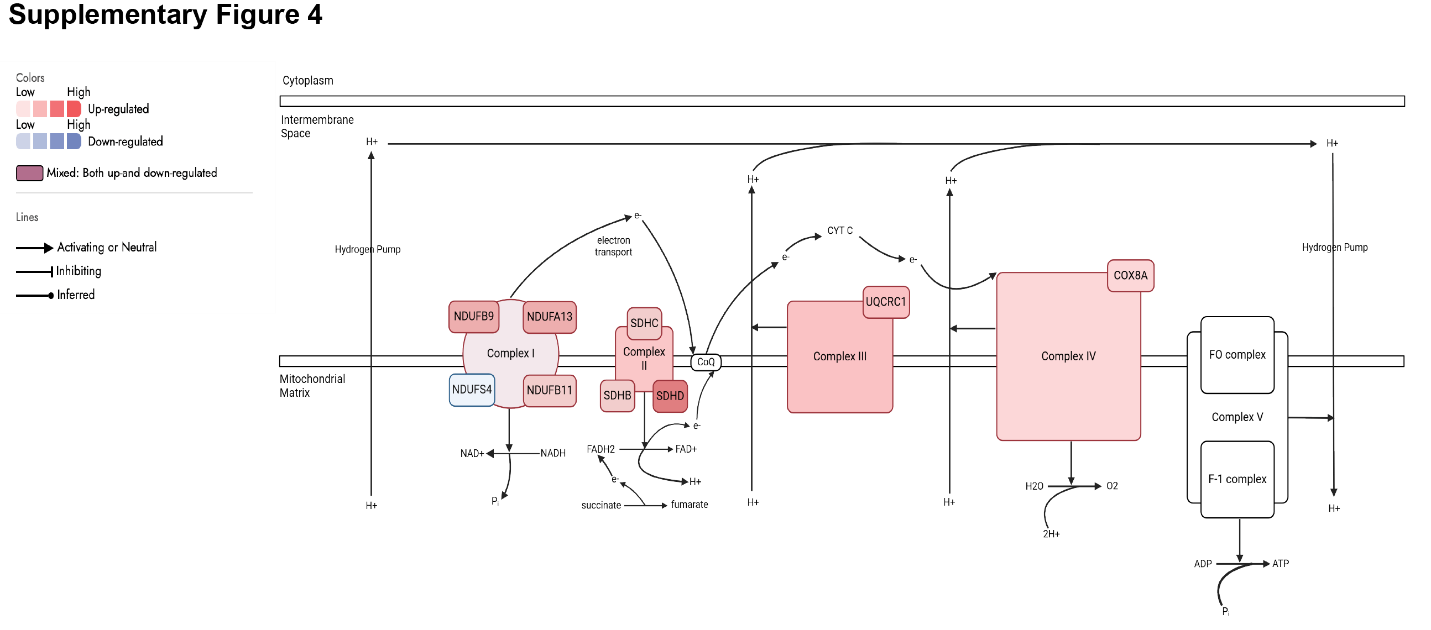

**Supplementary Figure 4. Increased expression of Complex II-IV subunits in the NDUFS4 KO NSCs.** Altered mitochondrial genes associated with respiratory chain complexes identified from DEGs of NDUFS4 KO NSCs. Figure drawn using BioRender.

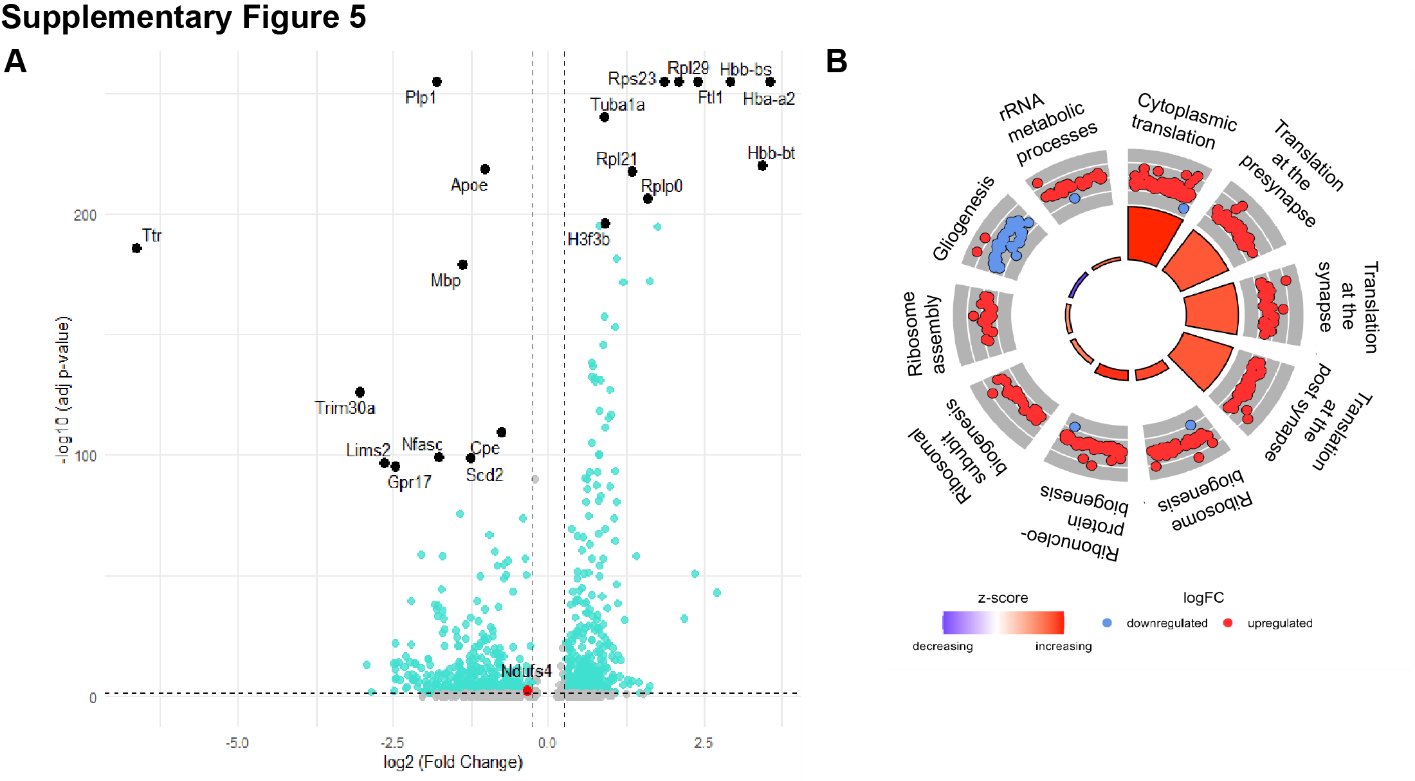

**Supplementary Figure 5. Upregulated protein translation pathways in NDUFS4 KO neuroblasts. A.** Volcano plot showing fold change of differentially expressed genes in NDUFS4 KO neuroblasts. Top 10 up and downregulated mRNAs are labelled (Black circles) and NDUFS4 is highlighted in the red circle. The threshold was set at log2(Fold Change) ≥ +0.25 and ≤ -0.25, and –log10 (adj p-value) < 1.2991 **B.** GO circle plot showing top dysregulated biological processes and regulation of associated genes in NDUFS4 KO neuroblasts.

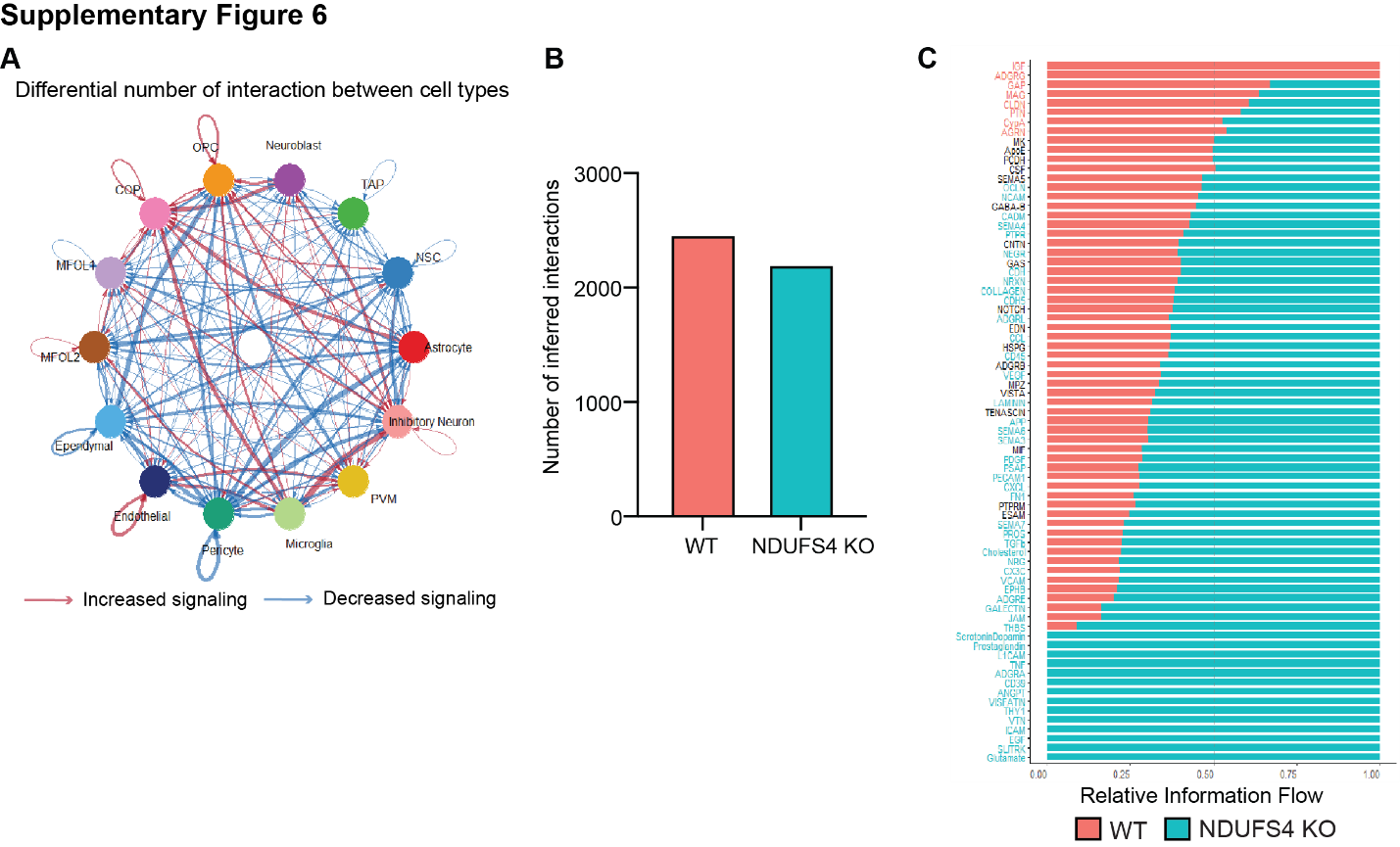
**Supplementary Figure 6. The number of signaling interactions is decreased in most cell types including neural progenitors in NDUFS4 KO as compared to WT.** **A.** Differential number of interactions between cell types from NDUFS4 KO. Blue and red lines indicate reduced and increased interactions between cells, respectively. **B.** Total number of signaling interactions in all cells. **C.** The significant signaling pathways were ranked based on their differences of overall information flow within the inferred networks between WT and NDUFS4 KO. The overall information flow of a signaling network is calculated by summarizing all the communication probabilities in that network. The top signaling pathways colored by red are more enriched in WT, and the bottom ones colored by cyan were more enriched in the NDUFS4 KO.

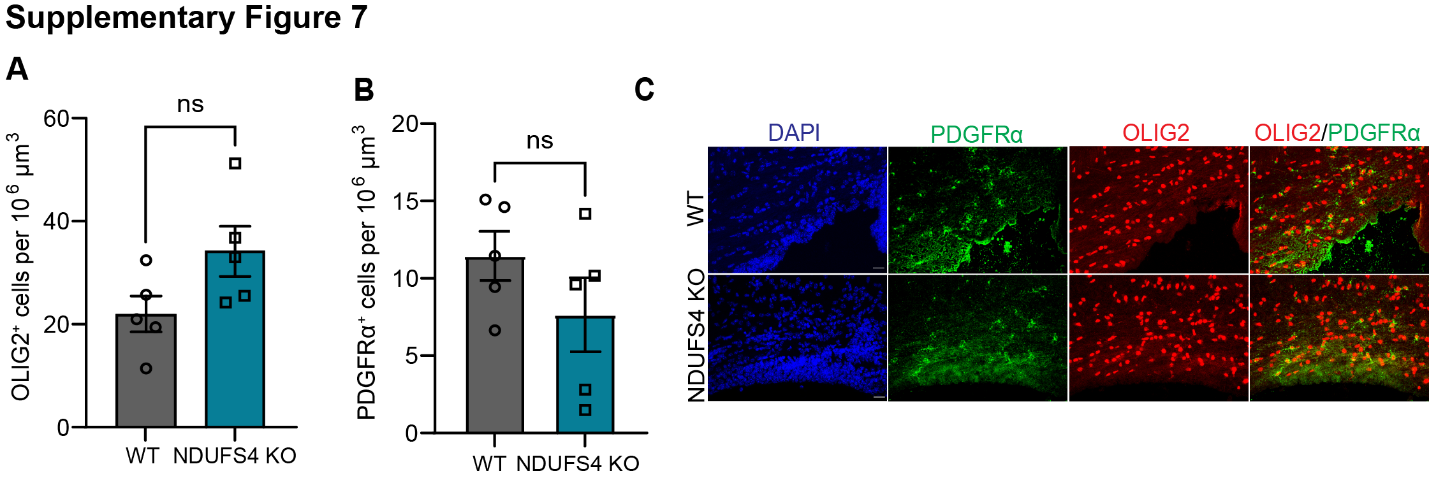

**Supplementary Figure 7: Oligodendrocytes are not changed in NDUFS4 KO SVZ at early postnatal day P14. A-B.** Density of (A) oligodendrocytes (OLIG2^+^) and (B) oligodendrocyte progenitors (PDGFRα^+^) at P14. **C.** Representative confocal images of immunohistochemical detection of PDGFRα (green), OLIG2 (red), and DAPI (blue). Scale bar= 20 um. ns = not significant. Each dot represents one animal.

**Table S1: Differentially expressed gene lists for neural stem cells comparing WT and NDUFS4 KO mice.**

**Table S2: Differentially expressed gene lists for TAPs comparing WT and NDUFS4 KO mice.**

**Table S3: Differentially expressed gene lists for neuroblasts comparing WT and NDUFS4 KO mice.**
